# Low-invasive, wide-field, and cellular resolution two-photon imaging of neural population activity in brainstem and nucleus tractus solitarii

**DOI:** 10.1101/2024.05.31.596631

**Authors:** Masakazu Agetsuma, Azumi Hatakeyama, Daisuke Yamada, Hiroshi Kuniishi, Eri Takeuchi, Shinji Tsuji, Chihiro Ito, Tomoko Kobayashi, Atsushi Noritake, Yoshitsugu Aoki, Hiroshi Yukawa, Akiyoshi Saitoh, Junichi Nabekura, Masayuki Sekiguchi

## Abstract

Brain-viscera communication plays a crucial role in regulating mental health, with the vagus nerve being a key structure mediating this interaction. Clinically, artificial vagus nerve stimulation (VNS) is used to treat various neuropsychiatric disorders, highlighting the importance of vagal afferent fibers in regulating emotion. The nucleus tractus solitarii (NTS) is a brainstem structure proposed to receive signals from vagal afferents and relay them to brain networks for emotion regulation. However, due to the anatomical complexity and difficulty in accessing the deep-brain NTS region in living animals, the mechanisms remain unclear. Here, we developed a wide-field and deep-brain two-photon imaging method using a double-prism based optical interface. This approach enables the identification of cellular-resolution neural activities in the NTS while preserving the cerebellum, which covers the NTS and is important for emotion regulation, intact. We systematically evaluated how NTS neurons respond to VNS and a gastrointestinal hormone, suggesting the usefulness of this method for investigating the role of the vagus-NTS pathway *in vivo*.

## Introduction

Optical techniques are powerful for measuring and artificially manipulating neuronal activity to elucidate the function of neuronal circuits and dynamic information processing in the brain. These techniques have been widely used in rodent brain research. However, technical limitations remain in accessing deep brain regions. For example, the brainstem, a key deep-brain region for brain-viscera communication, has been difficult to optically investigate its physiological properties.

Regulating emotion through brain-viscera communication has received significant attention, and the abnormal communication is linked to neuropsychiatric disorders^1–6^. Afferent vagus fibers originating in the nodose ganglion in the cervical cord play a crucial role in brain-viscera communication, receiving information from various internal organs and relaying it to the central nervous system through the brainstem area^5,6^. Artificial vagus nerve stimulation (VNS), which is clinically applied to treat various neurological and psychiatric disorders^7–9^, mainly target the afferent vagus nerve. Initially reported in the 19th century, the FDA approved VNS for treating drug-resistant seizures in 1997 and depression in 2005^10,11^. Currently, several neuropsychiatric and related disorders are considered potentially treatable with VNS^12^. For safe and effective VNS, the parameter settings must be carefully optimized^13,14^. Research using rodent models is crucial for this optimization by elucidating the mechanisms underlying VNS and its therapeutic effects. Previous studies in rodents demonstrated that VNS effectively suppresses seizures^15,16^, mitigates cerebral ischemia^17^, and treats chronic pain^18^. Furthermore, VNS significantly reduces depressive symptoms^19,20^ and facilitates the extinction of associative fear memory^21^. These findings suggest that the insights gained from animal models are critical to advancing VNS-based clinical treatments and understanding the mechanisms of emotion regulation through the vagus nerve.

Previous studies have indicated that the nucleus tractus solitarii (NTS) is likely a critical gateway for regulating emotions through brain-viscera communication and for VNS-based clinical treatments^6,8,22,23^. Afferent vagus fibers receive information from various internal organs and are proposed to relay it to the NTS^6,23–26^. The NTS projects to various brain regions involved in emotion regulation, such as the amygdala, lateral hypothalamic area, bed nucleus of the stria terminalis, and periaqueductal gray^6,8^, which suggest the involvement of the vagus-NTS pathway in regulating emotion. VNS has also been shown to modulate the activity of various brain regions^27–29^, including the locus coeruleus, whose activity is regulated by the NTS. Lesions of the locus coeruleus suppress the anti-epileptic effects of VNS^15^. The actual role of the NTS and the detailed mechanisms underlying the therapeutic effects of VNS, however, remain unclear. It is also unclear how information from various organs relayed to the NTS for regulating emotion is processed, and whether the information from various organs is segregated or integrated within the NTS.

Addressing these questions has been challenging because the NTS is located deep in the brain and is surrounded by regions critically involved in vital functions, making direct observation difficult. Removing the cerebellum to expose the brainstem allows for observations of a wide area of the brainstem including the NTS, which is useful to study the role of the NTS in regulating the information from the digestive systems^30^. However, the cerebellum is also an essential brain structure for emotional regulation^31–34^ and is modulated by the VNS-based treatment^28^. Therefore, the development of a novel technique for *in vivo* recordings of brainstem neurons without removing the cerebellum is in high demand. A technique to optically access the brainstem and NTS in the deep brain area while keeping the cerebellum intact would expand the possibilities for neural activity imaging and optogenetic neural activity manipulation, thereby profoundly advancing our understanding of the function of the NTS and its contribution to emotional regulation.

This aim motivated us to develop an *in vivo* two-photon imaging technique using a double-prism based optical interface (<u>D</u>ouble<u>-P</u>rism based brain<u>S</u>tem imaging under <u>C</u>erebellar <u>A</u>rchitecture and <u>N</u>eural circuits, "D-PSCAN"). The D-PSCAN method enables the observation of neural population activity with single-cell resolution in the brainstem and NTS situated deeply beneath the cerebellum while keeping the cerebellum and surrounding neural connections intact. We demonstrated the capability of this method to distinguish NTS activity from that of descending vagus neurons whose cell bodies lie ventrally to the NTS. The recorded neural activity data were used to evaluate the physiological characteristics of NTS neurons, i.e., their responsiveness to VNS. We also demonstrated the application of this imaging technique for detecting responsiveness to a gastrointestinal hormone cholecystokinin-8 (CCK), a more physiological stimulus than VNS. Our results suggest the applicability of this method for detailed understanding of the physiological mechanism of the NTS and future research on emotional regulation through the vagus-NTS pathway.

## Results

### Double microprism for optical access to the brainstem area deep below the cerebellum

We developed a novel *in vivo* two-photon calcium imaging method for observing the deep brain area in mice that enables neural population activity recordings in the brainstem and NTS. Using a genetically encoded Ca^2+^ indicator, GCaMP, expressed by an adeno-associated virus (AAV), we aimed to detect neural activities in the NTS (Figure 1A–B) to observe the responsiveness of NTS neurons to various stimuli. Viruses encoding GCaMP were injected into the NTS at approximately postnatal day (P) 50–60 for *in vivo* imaging experiments, at least 1 month before the microprism implantation described below. Although primarily targeted at the NTS, viruses often spread to adjacent areas, beneficially inducing additional expression in surrounding regions and allowing us to visualize wide areas of the brainstem.

**Figure 1.**
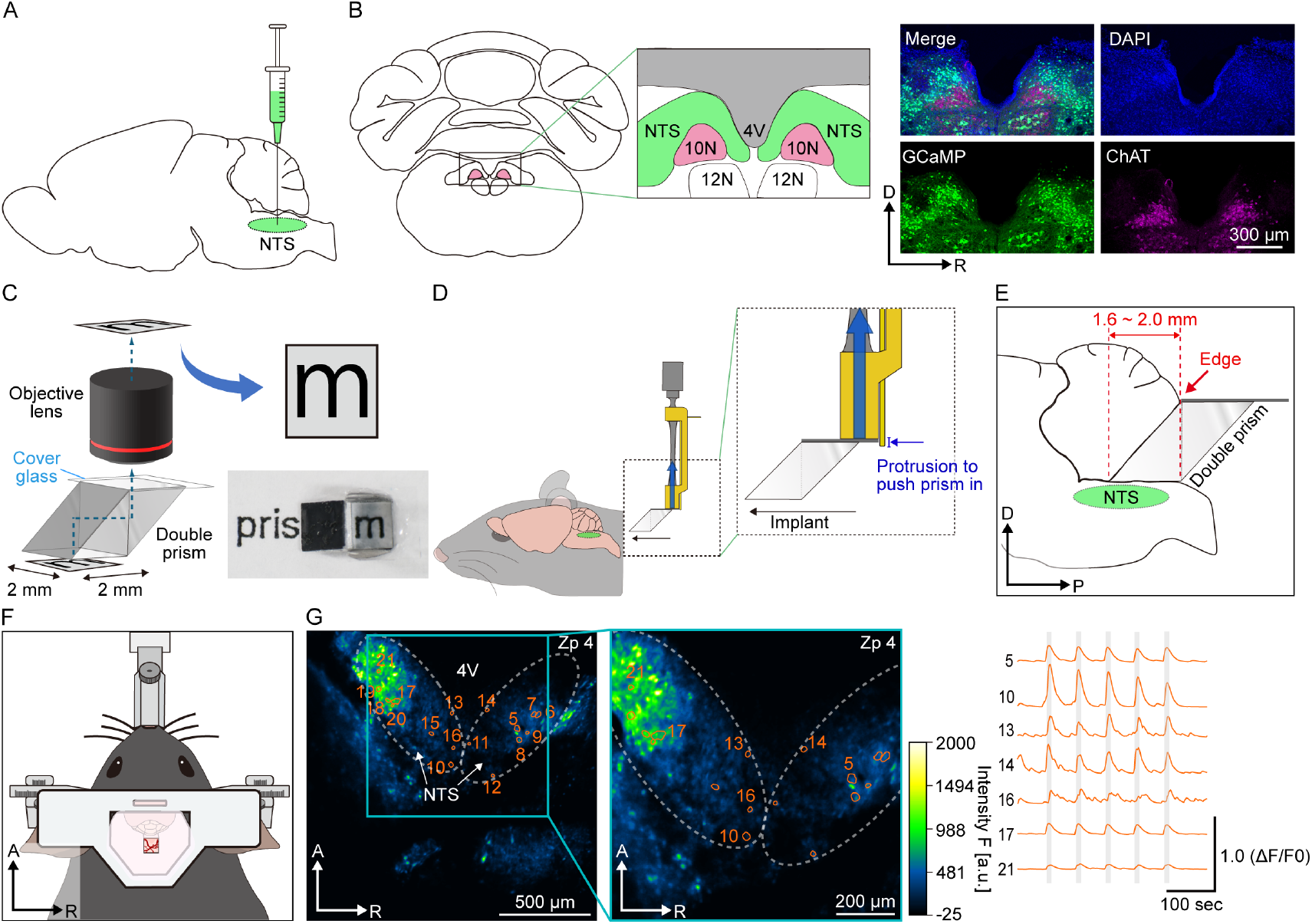
Schematic overview of *in vivo* NTS imaging with a double prism (A) Sagittal view of a mouse brain illustrating how GCaMP was expressed in NTS neurons by AAV injection.(B) GCaMP expression in the NTS (coronal view). (left) Schematic illustration of the injection site modified from the Allen Brain Atlas. (right) Expression of GCaMP (green) in the NTS, exclusive to the 10N area (marked by ChAT immunohistochemical staining in magenta). Blue, DAPI. (C) Two 2-mm right-angled prisms are attached to each other using UV-curable optical adhesive and bonded to a 3 mm × 3 mm square cover glass. In this illustration, the letter “m” at the bottom of the double prism appears through the double prism and the objective lens. Similarly, in the picture at the right bottom corner, the letter “m” in the word “prism” appears through the double prism. (D) A vacuum-type prism holder is used to implant the double prism. Black arrows indicate the directions in which the double prisms are pushed forward. Thick blue arrows indicate directions in which the double prisms are vacuumed up to be held by the holder. A thin blue arrow indicates the protrusion to push the double prism during the implantation. (E) Implantation of the double prism. The double prism is carefully inserted between the cerebellum and the brainstem approximately 1.6‒2.0 mm from the posterior edge of the cerebellum. The area containing the NTS is observed through the double prism. (F) A head plate is attached to the mouse skull using dental cement after the microprism implantation. (G) Example of *in vivo* brainstem imaging showing neural activity in a live mouse. Left: Representative two-photon images showing GCaMP fluorescence in a large field of view covering the wide brainstem. Each number in the image indicates a region of interest (ROI), each corresponding to a putative single neuron. Median fluorescence over time is presented in green-fire-blue to show a baseline fluorescence level. Middle: Magnified view of a section from the left panel. Right: Representative traces showing changes in GCaMP signal over baseline (ΔF/F0) in selected ROIs. Gray indicates the timing of VNS. Zp4, 4th z-stacked plane obtained by the volumetric scanning (see also Figure 2A and Figure S3). D, dorsal; R, right; P, posterior; A, anterior.

The brainstem is located deeply below (2.0–2.5 mm) the dorsal surface of the cerebellum. One potential problem in observing neural activities in the brainstem is that even when the mouse head is fixed under the objective lens for two-photon imaging by tightly attaching the head plate to the skull, the brainstem can still move somewhat independently in response to the mouse’s body movements^30^. Although standard skull reference points (bregma and lambda) are typically referred to when targeting specific brain areas, it is also crucial to visually capture the brain structure under observation in each mouse. In a previous study, the cerebellum was removed to address these issues^30^, but the cerebellum is also involved in emotion control^31^; therefore, to accurately elucidate the role of the vagus-NTS pathway in regulating emotion, it is essential to have a wide field of view of the brainstem while keeping the cerebellum and neural connections intact.

To achieve this aim, we developed a technique to optically access the deep brainstem area using a double prism (Figure 1C–E). We designed an implantable double microprism assembly comprising a 2-mm right-angle glass microprism bonded to another 2-mm right-angle glass microprism and a 4 mm × 4 mm or 3 mm × 4 mm cover glass window at the surface-end position of the imaging window (Figure 1C). Dental cement was used to fix this assembly at the correct position for NTS imaging. The cover glass enhanced adhesion to the dental cement, thereby securing the position of the double microprism assembly and preventing dental cement from obstructing its optical path. We implanted the assembly (simply termed "double prism") between the cerebellum and the brainstem using a custom-made vacuum-based manipulator (Figure 1D and Figure S1A–C). To implant the double prism, the skull over the dorsal surface of the cerebellum was carefully removed using a dental drill and fine forceps (Figure S1D–E). A micro-knife was used to incise the thick dura mater between the posterior edge of the cerebellum and the spinal cord. Removing the dura mater over the cerebellum not only allows for smooth insertion of the prism but also adequately relieves the pressure that might occur when the prism is inserted between the cerebellum and brainstem.

Next, the newly developed vacuum-based prism holder was used to insert and implant the double prism between the cerebellum and brainstem (Figure 1D–E, Figure S1C, F– G). Sufficient space and no neural fibers remained between the cerebellum and brainstem around the NTS area, enabling smooth insertion of the prism without cutting neural fibers. Using a stereotaxic frame and vacuum-based prism holder attached to an equipped manipulator, the prism can be precisely operated during the implantation while being maintained in a horizontal position. The vacuum-based prism holder has a hook on one side (Figure 1D and Figure S1B–C), which exerts a pushing force when sliding the sharp corner of the double prism between the cerebellum and brainstem from the caudal side to the rostral side (Figure 1D–E). After slowly inserting the double prism to approximately 1.6 mm deep, we waited a few minutes to ensure that breathing and other physical conditions were not disturbed (Figure 1E). The double prism was slid further down at 0.1-mm increments until reaching a maximum depth of 2.0 mm. Any physical reactions such as disrupted breathing occurring during the insertion led us to promptly retract the prism. Setting a stereomicroscope at an angle such that the brainstem can be visualized through the double prism allows for clear observation of the arrangement of blood vessels in the brainstem and fourth ventricle while inserting the prism.

After the double prism was fully inserted, UV craft resin was used to fix the double prism and coat the surface of the targeted nerve structures (Figure S1G), followed by tight fixation with dental cement. A resin coating was applied with the double prism remaining in the holder and maintaining sufficient contact with the brainstem target area (including the NTS) to prevent the resin from getting between them. After the UV resin had cured, the holder was carefully released, and the double prism was firmly fixed to the surrounding areas, including the remaining skull bone rostral to the cerebellum, using dental cement. For further head fixation for two-photon imaging under the microscope objective, the head plate was attached to the skull with dental cement (Figure 1F).

The double prism allows for optical access to the brainstem located on the ventral side of the cerebellum, which exceeds 2.0 mm in thickness. Using the wide field of view and horizontal coordinates, the position of the fourth ventricle serves as a key reference for locating the NTS. Two-photon imaging through the double prism enabled us to detect neural activities in the NTS at single-cell resolution, visualized by GCaMP (Figure 1G). We named this method "D-PSCAN" as explained above.

We confirmed the specificity of the method for detecting NTS neural activities. A primary purpose for developing this imaging method was to investigate how information from afferent vagus nerve inputs is processed in the NTS. On the other hand, the dorsal motor nucleus of the vagus nerve (10N, also known as DMNV or DMV), where the cell bodies of the vagal efferent fibers originate, is located just ventrally to the NTS^6^, suggesting the importance of confirming specificity in the NTS and distinguishing NTS neurons from 10N neurons. Importantly, the two-photon imaging technique offers superior spatial resolution at depth. Based on this advantage, we confirmed that the obtained signals originated from the brainstem surface area, i.e., the NTS, rather than deeper regions such as 10N. To further test the specificity, we prepared fixed brain slices after the *in vivo* imaging experiments and checked the expression of GCaMP within the NTS. We used ChAT, an immunohistochemical marker, to identify 10N neurons^35^ and distinguish NTS neurons histologically. Expression of GCaMP was not observed in ChAT-positive cells within 10N (Figure 1B), indicating that NTS neuron activities can be detected separately from efferent vagus neuron activities using this method. We also checked for unanticipated retrograde labeling of the vagal afferent terminals onto the NTS. Cell bodies of vagus neurons projecting to the NTS are located in the nodose ganglion^6,36^. In mice that showed GCaMP expression in the NTS neurons, we observed no GCaMP-expressing cell bodies in the nodose ganglion, i.e., vagus sensory neurons (Figure S2). These findings demonstrated that, when imaging neural activities around the NTS region using this method, we can distinguish activities of NTS neurons from the signals of vagal afferent nerve terminals. Thus, the D-PSCAN method enables us to specifically evaluate the activities of the NTS neurons separately from the activities of vagal efferent neurons and vagal afferent fiber terminals.

### *In vivo* imaging of neural population activities at single-cell resolution in the brainstem (NTS) of live mice

Using the D-PSCAN method, we performed *in vivo* two-photon imaging and recorded the neural population activities of the NTS and surrounding brainstem areas. Spontaneous activities and stimulus-evoked responses of the brainstem neurons were recorded by imaging GCaMP fluorescence changes (ΔF/F0). Brain motion artifacts interfered with imaging in awake mice but were minimized under anesthesia, enabling stable detection of the neural activities at a single-cell resolution both inside and outside of the NTS area (Figure 2 and Supplementary Movie 1, 2). Time-lapse imaging at various depths was performed by adjusting the imaging focal plane using a piezo to move up and down in depth (Figure 2A and Figure S3). Volumetric live imaging (2 mm x 2 mm horizontally, or smaller for zooming, 200–480 μm dorsoventrally, at 0.4–1.0 Hz temporal resolution) of neural activities in the wide brainstem area (including the NTS) of the head-fixed mouse was performed through the implanted double prism. Importantly, this wide field of view and piezo-based volumetric imaging allowed us to detect GCaMP fluorescence and neural activities not only from inside but also from outside of the NTS areas and to confirm the specificity of the stimulus-evoked responses in the NTS, as explained below.

**Figure 2.**
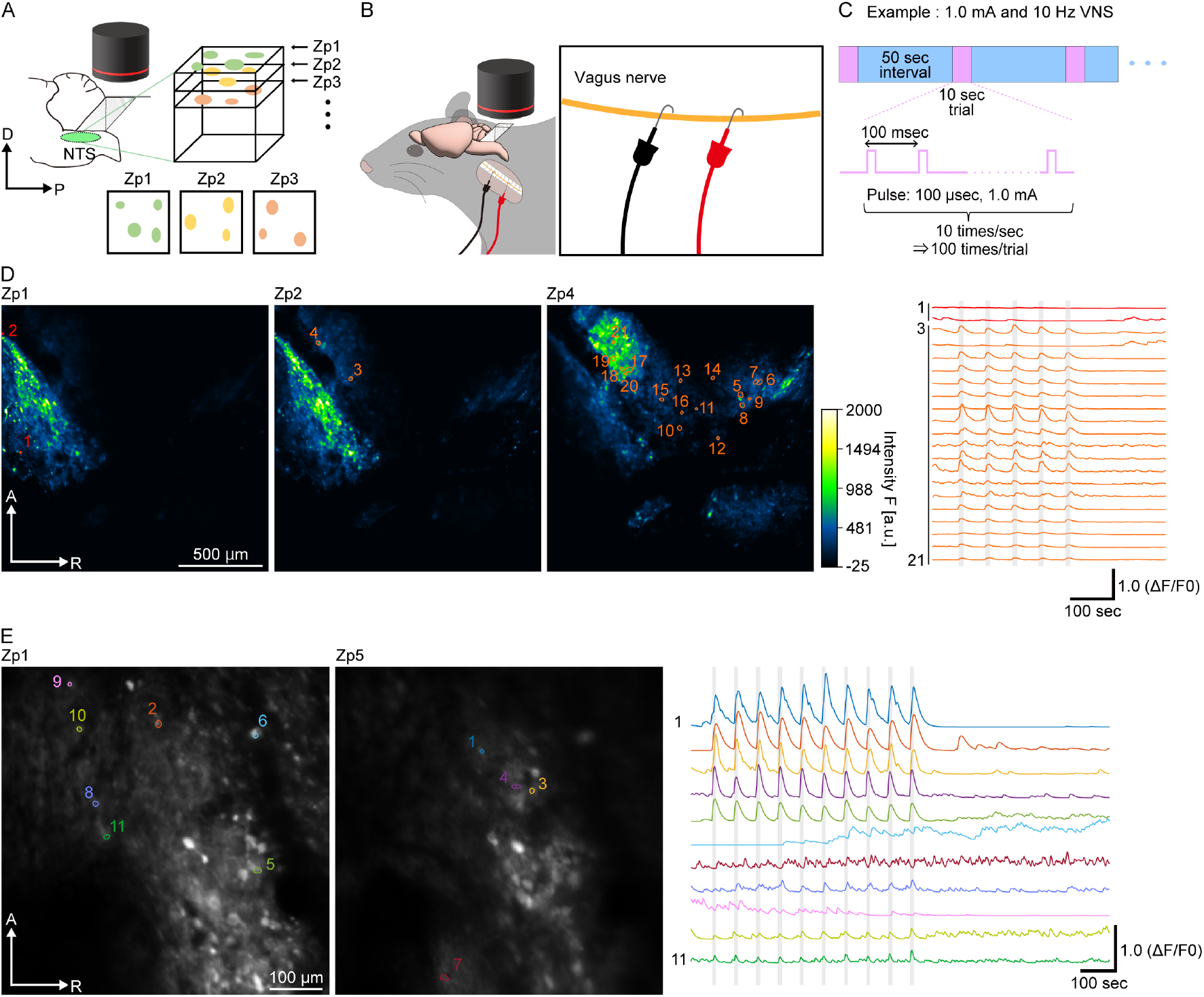
*In vivo* volumetric imaging of NTS activity during repeated VNS (A) Volumetric live imaging of neural activities in the wide brainstem area, including the NTS. Two-photon imaging was performed through the objective lens (black) and the implanted double prism in the head-fixed mouse under anesthesia. The wide field of view and piezo-based z-scanning enabled volumetric imaging of the GCaMP fluorescence. Colored circles illustrate examples of detected regions of interest (ROIs) at Zp1, Zp2, and Zp3. (B) *In vivo* NTS imaging was conducted during electrical stimulation of the vagus nerve through a bipolar platinum electrode, which wrapped the cervical vagal trunk in a mouse. The electrode was implanted and fixed with the insulating silicone adhesive. To create anodal block for efferent direction, cathode (black) and anode (red) were located on rostral and caudal site of vagus nerve, respectively. (C) Schematic of VNS protocol. In this example, a 1.0 mA, 10 Hz, 100 μsec train of electrical stimulation was delivered to the left vagus nerve via an electrode. The term "trial" refers to a VNS with a pulse width of 100 μsec for 10 sec, regardless of frequency and current intensity. The term "session" is defined as a series of 10-sec VNS presentations at a fixed frequency and current intensity, with a 50-sec interval, and this example represents a session at 1.0 mA and 10 Hz. (D) An example of *in vivo* volumetric imaging of the NTS during VNS at 2.0 mA and 10 Hz. AAVdj/Syn.GCaMP7f was used for this experiment. (Left) As examples, Zp1, 2, and 4, which contain automatically detected ROIs in this mouse, are shown (Zp4 here is the same data as Figure 1G). Median fluorescence over time for each panel is presented in green-fire-blue to show a baseline fluorescence level. (Right) Detection of spontaneous and/or VNS-evoked neural activities (ΔF/F0) from 21 example neurons outside (red) and inside (orange) the NTS. See also Supplementary Movie1. (E) Higher-magnification imaging provides clear single-cellular precision. NTS imaging was performed during VNS at 0.5 mA and 10 Hz. AAV1/Syn.GCaMP6f was used for this experiment. (Left) As examples, Zp1, and 5, which contain automatically detected ROIs in this mouse, are shown. To clearly distinguish the ROIs by color, median GCaMP fluorescence in the NTS is shown in grayscale. (Right) Eleven ROIs, each corresponding to a putative single neuron, are sorted by the magnitude of their responses to the VNS. See also Supplementary Movie2. Zp, z-stacked horizontal plane obtained by the volumetric scanning. Gray indicates the timing of VNS. D, dorsal; P, posterior; A, anterior; R, right.

### Detection of single cell activities in the NTS as the direct efferent target of the vagus nerve

To demonstrate the usefulness of the D-PSCAN for investigating the physiological characteristics of the NTS neurons, we applied the method to observe the neural responses to artificial electrical stimulation of the afferent vagus nerve under various conditions (vagus nerve stimulation, or VNS). We tested whether NTS neurons respond to the stimuli and further evaluated the diversity of their responsiveness. To perform VNS, the bipolar platinum hook electrode that has been used in rodents^37^ was employed (Figure 2B). After isolating the vagus nerve from the carotid artery, a bipolar platinum hook electrode was wrapped around the left vagus nerve and fixed in place without contacting other tissues using the insulating silicone adhesive (Kwik-Sil)^38,39^. The electrode was positioned to keep the positive and negative poles apart. This setup allows for stable activation of the vagus nerve while preventing irritation to the surrounding tissue. To create an anodal block for the efferent direction, the cathode and anode were located on the rostral and caudal sides, respectively, of the vagus nerve. The mechanism of the anodal block is proposed to selectively bias activation toward vagal afferent fibers^40–44^. We selectively targeted the left vagus nerve, which is usually chosen for safety reasons, as stimulation of the right vagus nerve has a greater impact on heart rate^45,46^. We conducted *in vivo* NTS imaging experiments after closing the surgical wound.

We imaged the *in vivo* neural activity in the brainstem as VNS was repeatedly applied. The VNS parameters were established based on previous studies, and various conditions (i.e., various current intensities and stimulus frequencies) within the ranges reported in prior rodent and human studies^13,42,43,47^ were used to investigate the neural response properties. Figure 2C presents an example in which the stimulation intensity was set to 1.0 mA and the stimulation frequency was 10 Hz. A pulse width of 100 microseconds (μsec) was consistently selected across all experiments in the present study. At a stimulus frequency of 10 Hz, stimulation (100-μsec current) occurred 10 times per second. The term “trial” is defined as each 10-sec VNS duration, while the term “session” is defined as a series of VNS trials at each condition (e.g., 1.0 mA at 10 Hz, 0.5 mA at 10 Hz, 1.0 mA at 5 Hz, etc.). In each session, the inter-trial interval was set at 50 sec. In the example cases shown in Figure 2, 10-sec VNS was repeated multiple times within a session to assess the neural responses to each VNS condition.

Taking advantage of the 2 mm x 2 mm large field of view provided by the double prism, we conducted volumetric observations of a wide brainstem area, including the NTS, and compared neural responses to VNS inside and outside of the NTS (Figure 2D). In an example case (Figure 2D), we scanned a total of 16 z-planes at each time point. We stacked every 4 consecutive plane from top to bottom to create an average image for each time point with carefully correcting the motion artifact (see Methods for details). This stacking was done with a two-plane overlap; for instance, the first stack included the 1st to 4th z-planes, the second stack included the 3rd to 6th z-planes, and subsequent stacks followed this pattern. As a result, we obtained 7 stacked planes in total in this experiment, labeled Zp1 through Zp7, respectively. This procedure is advantageous to enhance the signal to noise ratio in each movie, which is critical for further automatic detection of the active neurons^48^. The motion-corrected movies were analyzed to automatically detect the active neurons using a constrained nonnegative matrix factorization algorithm, as described previously^48,49^. This detection method is advantageous in isolating the activity of the individual neuron, even when other active neurons are nearby or overlapped within the field of view^48^ (see Methods for details). After automatic detection of the active neurons, we checked the existence of overlapped detection of the regions of interest (ROIs, i.e., active neurons) between adjacent Zps, and removed such overlaps by manual inspection.

Temporal dynamics of neural activity were clearly observed within the field of view at multiple depths (Figure 2D and Supplementary Movie 1). Here, we present the GCaMP signals at Zp1, Zp2, and Zp4. Neurons exhibiting stimulus-evoked and/or spontaneous neural activity were automatically identified, and representative neurons are shown in the graph. In the case of neurons outside of the NTS, we observed no clear responses to VNS and detected only spontaneous (stimulus-independent) activities (neurons #1 and #2 in Figure 2D). On the other hand, we detected a large number of neurons responding to VNS in the NTS area (neurons #3–#21).

Two-photon imaging utilizes a scanning-type observation system, enabling seamless transition from wide-field to high-magnification imaging using a single objective lens. Depending on the point spread function of the objective lens, this approach facilitates the acquisition of images with enhanced spatial resolution compared with maximum wide-field imaging. Using this mechanism, after confirming the position of the NTS in the wide field of view, we narrowed down the imaging area to the NTS and observed the neural activities at high magnification (Figure 2E and Supplementary Movie 2). We detected individual neuron cell bodies more clearly through visual inspection at higher magnifications compared with lower magnifications, confirming that most neurons identified at lower magnification were indeed NTS neuron cell bodies, which exhibited clear responses to VNS.

Note that there was variability in VNS responsiveness among neurons detected within the NTS (Figure 2D and E), prompting us to further examine their characteristics more thoroughly by varying the stimulus conditions.

### Diverse neuronal responses to VNS in the NTS

*In vivo* brainstem imaging was performed during VNS administered at various intensities and frequencies to explore how NTS neurons respond to different intensities and frequencies. Additionally, we assessed the responsiveness of the NTS neurons to repetitive stimulation across different intensities and frequencies.

First, we set the stimulus frequency at 10 Hz and varied the stimulus intensities (Figure 3A–E and Figures S4A, S5). Most of the stimulus-responsive NTS neurons exhibited enhanced responses to higher stimulus intensities. On the other hand, one neuron detected outside of the NTS showed only a spontaneous activity change and no clear response to VNS (neuron #2 in Figure 3 and Figures S4, S5). NTS neurons showed minimal responses to VNS at 0.1 mA and 0.25 mA and there was no clear difference between these intensities, while there was a substantial increase in the neural response at 0.5 mA. The difference between 0.5 mA and lower intensities suggests a possible threshold between them for recorded NTS neurons. At 1.0 mA, the responses increased further. We also calculated various parameters for evaluating the neural responses and observed consistent tendencies (Figure 3E).

**Figure 3.**
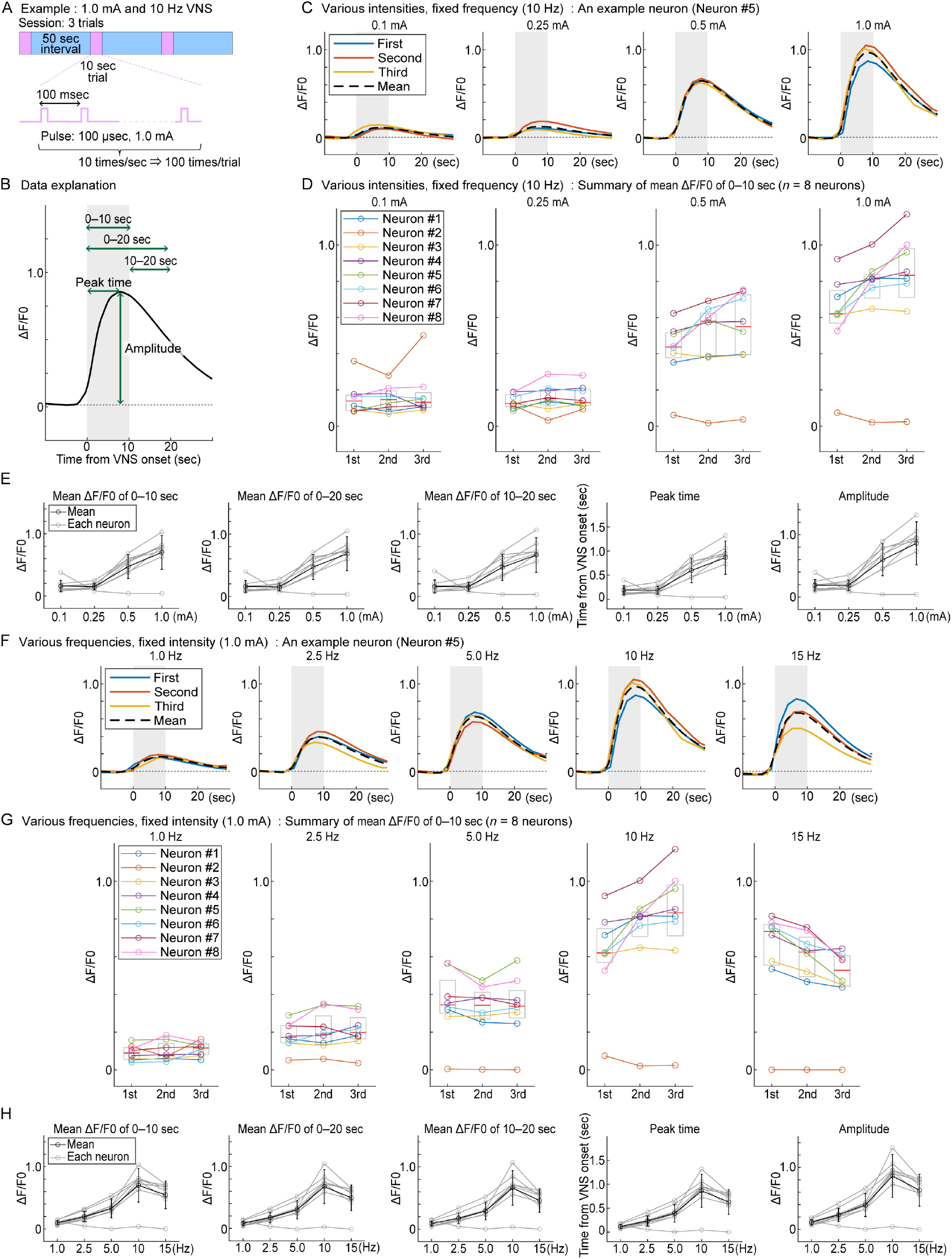
Variable responsiveness of NTS neurons characterized by various VNS conditions (A) Schematic of VNS protocol. In this example, a 1.0 mA, 10 Hz, 100 μsec train of electrical stimulation was delivered. Each experiment in this figure includes a session consisting of three trials of 10-sec VNS presentations, with a 50-sec interval. (B) Various parameters for evaluating neural responses. The neural response (ΔF/F0) to VNS (gray shaded area), with the horizontal axis representing the time from the VNS onset. Average signal (0‒10, 0‒20, 10‒20 sec), peak time, and amplitude, which are calculated for evaluating neural responses are indicated by green arrows. (C) Neural responses of an example neuron to VNS at a fixed frequency of 10 Hz, with various stimulus intensities. Colored lines indicate the neural response to the first, second, and third VNS. Dashed line indicates the mean of the responses to three VNSs. (D) Summary of responses of simultaneously recorded neurons (*n* = 8 neurons in a mouse) to VNS at a fixed frequency of 10 Hz with various stimulus intensities. The mean ΔF/F0 of each neuron for 10 sec from the onset of the VNS at various stimulus intensities are summarized. (E) Summary of neuronal responses to VNS at a fixed frequency of 10 Hz, with various stimulus intensities. Mean values for three trials, for each neuron (gray line) or for all neurons (black line), were calculated. Mean ΔF/F0 of 0–10 sec: the mean of ΔF/F0 for 10 sec from the onset of the VNS. Mean ΔF/F0 of 0‒20 sec: the mean of ΔF/F0 for 20 sec from the onset of the VNS. Mean ΔF/F0 of 10–20 sec: the mean of ΔF/F0 for 10 sec from the offset of the VNS. Peak time: time from the VNS onset to the ΔF/F0 peak. Amplitude: ΔF/F0 peak amplitude at each VNS presentation. (F) An example of ΔF/F0 of neuronal responses to VNS at a fixed stimulus intensity of 1.0 mA, with various frequencies. Colored lines indicate the neural response to the first, second, and third VNS. Dashed line indicates mean of the responses to three VNSs. (G) Summary of simultaneously recorded neurons (n = 8 neurons in a mouse) to VNS at a fixed stimulus intensity of 1.0 mA, with various frequencies. The mean ΔF/F0 of each neuron for 10 sec from the onset of the VNS at various stimulus frequencies are summarized. (H) Summary of neuronal responses to VNS at a fixed stimulus intensity of 1.0 mA, with various frequencies. Mean values for three trials, for each neuron (gray line) or for all neurons (black line), were calculated. Mean ΔF/F0 of 0–10 sec, mean ΔF/F0 of 0–20 sec, mean ΔF/F0 of 10– 20 sec, peak time, and amplitude were calculated as described in the legend of E. AAV1/Syn.GCaMP6s was used for this experiment. Gray shaded area indicates the timing of VNS. Boxes in D and G, 25th–75th percentile; Red bars in D and G, median of all neurons. Error bars in E and H, standard deviation.

We next evaluated the effect of repetitive stimulation on the neural responses at various stimulus intensities (Figure 3C–D and Figure S4A). At 0.1 mA and 0.25 mA, we did not observe a change in the responses to repeated stimulation (50-sec intervals) at each stimulus intensity. On the other hand, at higher intensities of 0.5 mA and 1.0 mA, some neurons exhibited gradually enhanced responses during the session with repeated stimulations.

Next, using the same recorded neurons, we set the stimulus intensity at 1.0 mA and varied the stimulus frequencies (Figure 3F–H and Figure S4B). The neuronal responses to VNS increased gradually as the stimulus frequency was increased from 1 Hz to 10 Hz, similar to when the stimulus intensity was increased. Interestingly, increasing the frequency to 15 Hz resulted in a reduction in the neural response compared with that at 10 Hz. We also investigated the effects of repetitive stimulation during each session at various frequencies. At 1 Hz, 2.5 Hz, and 5 Hz, we did not observe clear changes in response to repeated stimulation during each session. At 10 Hz, however, most of the NTS neurons showed a gradually increased response. Notably, at a higher frequency of 15 Hz, despite similar initial responses to those at 10 Hz, neurons exhibited decreased responses to subsequent stimuli during the session.

These results indicate that repeated stimulation may uniquely affect synaptic interactions and signal transmission between the vagal afferent nerve and NTS neurons. While we consider the physiological mechanisms behind the observed acceleration and deceleration below in the discussion section, our findings also suggest that the frequency-dependent increase and decrease in responses to VNS are not merely artifacts caused by the slow off-kinetics of GCaMP. Changes in neural activities and associated calcium concentrations occur more rapidly than the signal changes detected by calcium indicators like GCaMP. This discrepancy can theoretically lead to an artificial amplification of baseline signals during repetitive stimuli, which may account for the increase in fluorescent signals at 10 Hz. Our imaging system, however, reliably detected an opposing decrease in the GCaMP signal in response to repetitive stimulation at 15 Hz, despite similar initial response levels to those observed at 10 Hz. Thus, the enhanced responses at 10 Hz are also likely to reflect the actual responses of the neurons, rather than being artifacts.

Together, these findings indicate that the D-PSCAN method effectively detects various types of neurons in the NTS under different stimulus conditions. This method demonstrated that specific VNS conditions can directly and effectively modulate NTS neuron activities, potentially influencing whole-brain neural networks involved in the regulation of emotion.

### Chemical stimulation triggered both immediate and long-lasting responses in NTS neurons

Up to this point, we examined the response of NTS neurons to artificial electrical stimulation of the vagal afferent nerve. Emotional responses are also governed by more physiological stimuli and conditions. To assess the suitability of the D-PSCAN method for capturing responses under physiological conditions, we measured the activity of NTS neurons following intravenous administration of CCK (Figure 4A). CCK is a hormone released from the gastrointestinal tract after meals^50^. It directly influences the afferent fibers of the vagus nerve^51^ and is proposed to transmit signals via the NTS^51,52^ to various brain regions, including the feeding center.

**Figure 4.**
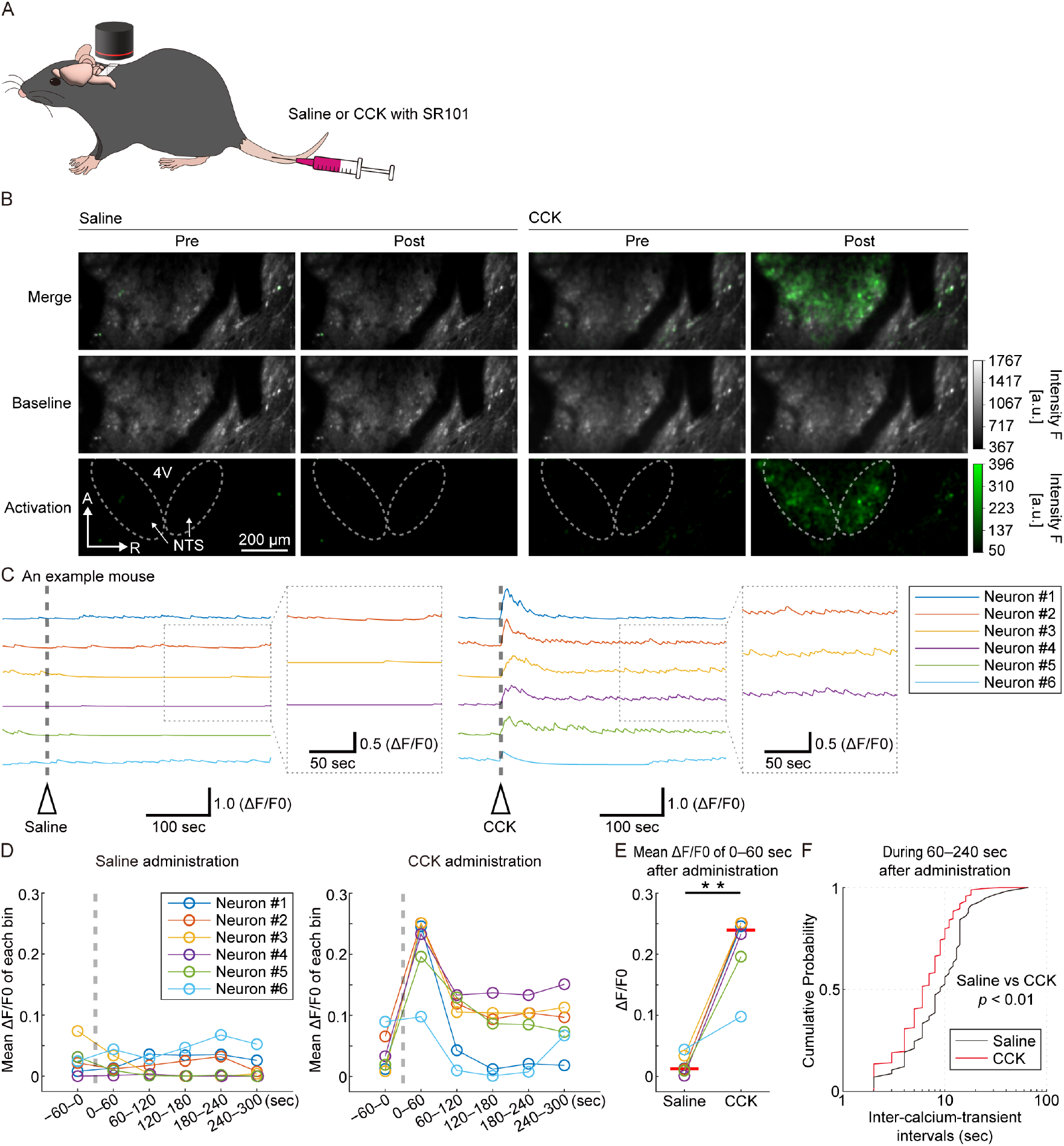
Imaging NTS neural activity in response to intravenous CCK administration (A) Schematic of *in vivo* NTS imaging while Saline or CCK was administered into the tail vein of an anesthetized mouse (co-administered with red fluorescent dye, SR101). Note that CCK administration was performed more than 10 min after saline administration. (B) Averaged fluorescence of GCaMP signals during pre- and post-administration (60 sec duration) of Saline or CCK. (Top) Baseline-subtracted signal is shown in green over a background of the baseline image shown in gray, (middle) baseline fluorescence image, (bottom) GCaMP signal. Dashed circles indicates the NTS area. 4V, fourth ventricle. See also Supplementary Movies 4 and 5. (C) Traces of GCaMP fluorescence in simultaneously recorded NTS neurons (same neurons on the left and right). The timings of the injections (saline, left; CCK, right) are indicated by a gray dotted line. Traces regarding more than five minutes after the injection are expanded to visualize the higher frequency of spontaneous activity (i.e., frequent calcium transients) uniquely observed after CCK administration. (D) The mean ΔF/F0 for each 60-sec bin along the time course was calculated for saline (left) or CCK (right) administration. Each data point represents the mean ΔF/F for each neuron in each bin. (E) The mean ΔF/F for 60 sec immediately after the administration of saline or CCK. Horizontal red bars, median of all neurons at each condition (saline or CCK). \*\**p* < 0.01, two-tailed Wilcoxon signed-rank test (*p* = 0.0011). (F) The cumulative distribution of intervals between calcium transient events from 60 to 240 seconds after the administration of saline or CCK. Neural activation was evaluated by detecting the timing of the calcium transients as described in Figure S6 (see also the Methods for more details). Data from all neurons (6 neurons shown in C) were pooled for each group for statistical comparison. \*\**p* < 0.01, Kolmogorov-Smirnov test (*p* = 0.0075). AAVdj/Syn.GCaMP7f was used for this experiment.

To assess whether the intravenous administration process itself affects neural activity, we also monitored responses during saline administration as a negative control. The timing of the intravenous administration was determined by including SR101 in the injected solution and observing the signal detected in the blood vessels surrounding the NTS using *in vivo* two-photon microscopy (Supplementary Movie 3).

With this method, we first demonstrated that saline administration did not induce a clear change in neural activity (Figure 4B–C and Supplementary Movie 4). In contrast, CCK administration induced a change in NTS neuron activities (Figure 4B–C and Supplementary Movie 5); upon administering CCK, we immediately observed clear and significant signal changes in NTS neurons (Figure 4B–E). This immediate response of the NTS neuron is consistent with a previous study demonstrating the responsiveness of vagal afferents to CCK^53^. Additionally, for more than 5 minutes after CCK administration, we observed a higher frequency of spontaneous activity compared with before CCK administration or after saline administration (Figure 4C, F and Figure S6). This prolonged activation in the NTS neuron is also consistent with the previous observation in vagal afferent fibers^53^. These results demonstrate that the D-PSCAN method is applicable to the detection of changes in NTS neuron activity induced under physiological conditions.

## Discussion

In the present study, to understand how the information processing mechanisms of brain-viscera pathways mediated by the NTS regulate emotions, we developed a new imaging technique named D-PSCAN based on *in vivo* two-photon imaging and double prism-based optical access. This method enables wide-field observation of deep-brain areas in the brainstem, including the NTS, while keeping the cerebellum, an important region for emotional regulation, intact. We confirmed that the D-PSCAN method allows for specific identification of NTS neurons, distinguishing them from surrounding regions based on horizontal coordinates in its wide field of view. Additionally, based on the advantages of two-photon imaging, we confirmed that the obtained signals originated from the surface brainstem area, i.e., the NTS, rather than deeper regions such as 10N. As we observed no GCaMP expression in 10N, it appears unlikely that vagal afferent terminals were labeled by GCaMP, ensuring almost no or undetectable contamination of their signal in the observed NTS areas. These findings together indicate that our method provides specific identification of NTS activities and assesses their responsiveness to VNS and CCK.

Using this method, we observed the responses of mouse NTS neurons to VNS, a technique clinically employed to treat neuropsychiatric disorders in humans. We confirmed that responses to electrical stimulation depended on the intensity and frequency, observed a threshold-like effect, and found that repeated excessive stimulation could reduce the responses. In addition to responses to artificial stimuli like VNS, we detected significant immediate responses and smaller but sustained activities following physiological stimuli such as CCK. These results demonstrate the usefulness of the D-PSCAN method for investigating how the NTS neurons process information from visceral organs for emotional regulation.

Recent studies have advanced our understanding of the complex mechanisms by which the NTS processes various types of visceral information^22,23,30,36,54^. How these mechanisms underlie the therapeutic effects of VNS and regulate emotion, however, remains unclear. Our imaging technique, which allows for detailed observation of NTS neuron activities while maintaining the integrity of cerebellar circuits crucial for emotional regulation^31–34^, is expected to be instrumental in deepening our knowledge of these complex mechanisms.

Repeated VNS induced both enhanced and diminished NTS responses, depending on the stimulus conditions. As discussed above, the observed NTS responses are unlikely to be artifacts caused by the calcium indicator (GCaMP). These results indicate that repeated stimulation may uniquely affect synaptic interactions and signal transmission between the vagal afferent nerve and NTS neurons. Inhibitory interneurons in the NTS are involved in information processing during mechanical stimulation of digestive organs^30,55^, suggesting the possible contribution of such complex circuitries within the NTS for adjusting the VNS signal. The activity of NTS neurons during VNS might also be modulated through interactions with other brain regions, including supramedullary regions^56^. Therefore, our imaging method will be useful for future investigations into these detailed regulatory mechanisms of the NTS. While an anodal block is likely to enhance specificity in targeting vagal afferent fibers, VNS might also weakly stimulate efferent fibers, which in turn might stimulate visceral organs and further activate the vagal afferent fibers. This may also influence the rates of the neuron responses. Overall, as optimizing VNS parameters for human patients is critical^13,14^, our NTS imaging method in mice could help reveal the underlying mechanisms of VNS, leading to more effective and personalized therapy settings.

Previous electrophysiological studies suggest that CCK induces prolonged activation of the vagal afferent nerve^53,57^. The CCK-evoked enhancement of the spontaneous fluctuations in the GCaMP signal lasting for more than 5 minutes is consistent with these electrophysiological observations. These results demonstrate that the D-PSCAN method has significant potential to detect changes in NTS neuron activity under physiological conditions.

Overall, our development of the wide-field and deep-brain two-photon imaging method utilizing a prism-based optical interface marks a significant advancement in the ability to study the NTS *in vivo* while maintaining an intact cerebellum. Future applications of this imaging system, particularly in awake animals, will be crucial for understanding the dynamic interactions within the NTS and its role in neuropsychiatric disorders. By enabling detailed observation of NTS activities, this method offers a powerful tool for advancing our understanding of the vagus nerve’s impact on brain function and emotional health, potentially leading to improved clinical interventions for neuropsychiatric conditions.

## Acknowledgements

We thank Okito Hashimoto for technical advice and discussion. We also thank the members of the laboratory for their help, especially Ikuko Takeda, Yuki Ichihashi, Ryo Fujikawa, and Fumiko Kito for experiments and data analyses. This study was supported by the JSPS KAKENHI Grant (grant number JP18K06536, JP20H05076, JP21H02801, JP22H05081, JP22H05519, JP24H01260 to M.A.), the Cooperative Study Program of National Institute for Physiological Sciences (19-233, 20-140 to M.S.; 21-244, 22NIPS218 to D.Y.), the practical research project for rare/intractable diseases from Agency for Medical Research and Development (AMED; Grant #: 20ek0109315h003 to M.S.), and the Research Grant for Neurological and Psychiatric Disorders from National Center of Neurology and Psychiatry, Japan (NCNP; Grant #: 30-3, 1-1, and 2-6 to M.S.).

## Author Contributions

M.A., D.Y. and M.S. conceived and coordinated the whole project. M.A. designed and developed a method for double-prism-based *in vivo* two photon imaging with the support of T.K. and J.N.; M.A., D.Y., H.K., E.T. and M.S. performed imaging experiments during the VNS and CCK-administration; M.A., A.H., D.Y., and S.T. performed data analyses with the support of C.I., A.N., H.Y., A.S. and M.S.; M.A., A.H., D.Y., A.N., Y.A., and M.S. wrote the paper, with contributions from all authors.

## Data availability

The data that support the findings of this study are available from the corresponding author upon reasonable request. The anatomical information in the Allen Brain Atlas (https://atlas.brain-map.org/) was used for the anatomical description, determination of the virus injection area, and evaluation of the recorded brain regions.

## Methods

### Animals

All animal experiments were carried out in accordance with National Institutes of Health guidelines and approved by the National Institute for Physiological Sciences Animal Care and Use Committee (approval number 18A102), or in accordance with the National Institute of Neuroscience, National Center of Neurology and Psychiatry (Tokyo, Japan) and were approved by the Institutional Animal Investigation Committee (approval number: 2020004). Male C57BL/6 mice housed under a 12-h light/dark cycle with free access to food and water were used for all experiments. Mice at 3–10 months of age were used for the experiments.

### Virus injection

Nucleus tractus solitarii (NTS) is a structure distributing in the dorsal medulla near the area postrema. To express GCaMP, a genetically encoded calcium indicator for monitoring neural activity, we used a gene expression system based on the AAV vector. We tested AAV1/Syn.GCaMP6s, AAV1/Syn.GCaMP6f, and AAVdj/Syn.GCaMP7f, which were produced as described previously^58,59^ using plasmids obtained from Addgene. As the GCaMP was expressed under the regulation of the Syn (Synapsin) promoter, all of the recording targets were assumed to be neuron-specific. All of the viruses allowed us to visualize the neural activity in the NTS, and type of virus used in each experiment is indicated in each corresponding figure legend. Viruses were injected into mice at approximately postnatal day (P) 50–60 for experiments at least 1 month before the microprism implantation, which was followed by *in vivo* imaging. Injection procedures were performed as described previously^60^, with some modifications to target the NTS efficiently. During surgery, the mice were anesthetized with isoflurane (initially 2% [partial pressure in air] and then reduced to 1%). A small circle (∼1 mm in diameter) of the skull was thinned over the cerebellum (around the injection site) using a dental drill to mark the site for a small craniotomy. Viruses were injected into the bilateral NTS regions at two sites (depth 4.7 and 3.7 mm from the dorsal surface of the cerebellum, volume 250 nl/site, at 100 nl/min) to sufficiently cover the NTS, over a 5-min period at each depth using a UMP3 microsyringe pump (World Precision Instruments). The X-Y coordinates for the injection site were usually 0.5 mm lateral to the midline and 3.3 mm caudal to lambda. The beveled side of the injection needle was faced to the caudal so that the needle could be smoothly inserted into and passed through the cerebellum, and the AAV would cover the NTS similarly in both hemispheres. We designed our injection protocol (especially the volume and depth) carefully to widely cover the area around the NTS, while the anatomical coordinates of the field of view for the two-photon imaging were precisely targeted using visual guide through the microprism implanted as described below (see the section "*In vivo* two-photon imaging in the wide brainstem area"). Also, as shown in Figure 1B, the expression of GCaMP was excluded in the 10N area, which placed next to and ventrally to the NTS as marked by the ChAT immunostaining. The reason for the exclusion is unclear, but it may be because the promoters and/or serotypes used in the present studies do not match for inducing the GCaMP expression in these ChAT-positive neurons. In any case, since neurons in the 10N area receive synaptic inputs from NTS neurons^6,61^ and are a source of vagal efferents, the lack of expression in the neurons of this area rather positively aids in the interpretation of the GCaMP signals detected by *in vivo* NTS imaging. In some mice, a deeper area (more than 0.5 mm from the dorsal surface of the brainstem, i.e., the surface of the microprism) also expressed the GCaMP when the fluorescence was checked in the brain slices (Figure 1B), but this deeper area was beyond the deepest limit of our two-photon imaging system for the detection of the sufficient signal intensity through the double prism *in vivo*. These results indicates that our recording system simply allows specific targeting of the neurons within the NTS and together with laterally-surrounding wide areas.

### *In vivo* two-photon imaging in the wide brainstem area

We developed a novel, minimally invasive method based on the double prism, which allows *in vivo* two-photon imaging of deep brain regions, specifically around the NTS. This innovative approach also enables simultaneous vagus nerve stimulation (VNS) or intravenous administration of cholecystokinin-8 (CCK).

*In vivo* two-photon imaging was performed as described previously^49,60^, with some special development for the optical access to the brainstem and NTS as explained below. Before the surgery, we designed an implantable double microprism assembly, comprising a 2-mm right-angle glass microprism (TS N-BK7, 2 mm AL+MgF2, Edmund) bonded to another 2-mm right-angle glass microprism and a 3 mm x 3 mm, 3 mm × 4 mm or 4 mm × 4 mm cover glass window (No.1; Matsunami) at the surface-end position of the imaging window, was prepared. Dental cement was used to fix this assembly at the correct position for NTS imaging. The cover glass was used to enhance adhesion to the dental cement, securing the position of the double microprism assembly and preventing dental cement from obstructing its optical path. We implanted the assembly (or simply termed "double prism") between the cerebellum and the brainstem using a custom-made vacuum-based manipulator (Figure 1D and Figures S1A–C).

At 1–8 months after the virus injection, the mice were anesthetized with isoflurane (initially 2% [partial pressure in air] and reduced to 1% during the surgery). For the implantation of the double prism, the skull over the dorsal surface of the cerebellum was carefully removed using a dental drill and fine forceps (Figure S1D–E). A micro-knife was used to incise the thick dura mater between the posterior edge of the cerebellum and the spinal cord. Removing the dura mater over the cerebellum not only allows for smooth insertion of the prism but also adequately relieves the pressure that might occur when the prism is inserted between the cerebellum and the brainstem.

Next, the newly developed vacuum-based prism holder was used to insert and implant the double prism between the cerebellum and the brainstem (Figure 1D–E, Figure S1B– C, F). Sufficient space and no neural fibers remained between the cerebellum and the brainstem around the NTS area, enabling smooth insertion of the prism without cutting neural fibers. Using a stereotaxic frame and vacuum-based prism holder attached to an equipped manipulator, the prism can be precisely operated during the implantation while being maintained in a horizontal position. The vacuum-based prism holder has a hook on one side (Figure 1D and Figure S1B–C), which exerts a pushing force when sliding the sharp corner of the double prism between the cerebellum and the brainstem from the caudal side to the rostral side (Figure 1D–E). After slowly inserting the double prism to approximately 1.6 mm deep, we waited a few minutes to ensure that breathing and other physical conditions were not disturbed (Figure 1E). The double prism was slid further down at 0.1-mm increments until reaching a maximum depth of 2.0 mm. Any physical reactions such as disrupted breathing occurring during the insertion led us to promptly retract the prism. When retracting the double prism assembly, it is crucial to ensure that the suction force by the vacuum is sufficient, as insufficient suction may lead to failure of the retraction. The area directly beneath the microprism was minimally compressed but remained intact, minimizing motion artifacts caused by the animal’s breathing. Setting a stereomicroscope at an angle such that the brainstem can be visualized through the double prism allows for clear observation of the arrangement of blood vessels in the brainstem and fourth ventricle while inserting the prism. Bleeding was rarely observed during the implantation procedure.

After the double prism was fully inserted, UV craft resin was used to fix the double prism and coat the surface of the targeted nerve structures (Figure S1G), followed by tight fixation with dental cement. A resin coating was applied with the double prism remaining in the holder and maintaining sufficient contact with the brainstem target area (including the NTS) to prevent the resin from getting between them. After the UV resin had cured, the holder was carefully released, and the double prism was firmly fixed to the surrounding areas, including the remaining skull bone rostral to the cerebellum, using dental cement. For the further head fixation for two-photon imaging under the microscope objective, the head plate was attached to the skull with dental cement (Figure 1F).

The double prism allows for optical access to the brainstem located on the ventral side of the cerebellum, which exceeds 2.0 mm in thickness. Using the wide field of view and horizontal coordinates, the position of the fourth ventricle serves as a key reference for locating the NTS. Two-photon imaging through the double prism enabled us to detect neural activities in the NTS at the single-cell resolution, visualized by GCaMP (Figure 1G). We named this method "D-PSCAN", or <u>D</u>ouble<u>-P</u>rism based brain<u>S</u>tem imaging under <u>C</u>erebellar <u>A</u>rchitecture and <u>N</u>eural circuits.

During the consecutive imaging sessions, mice were anesthetized with isoflurane (1%) or a ketamine (70 mg/kg) and xylazine (10.5 mg/kg) mixture (see the sections below for more details). Experiments were performed in a completely dark environment. The activity of NTS neurons was recorded by imaging fluorescence changes with a multiphoton microscope (Nikon A1R MP, Nikon) and a Mai Tai DeepSee Ti:sapphire laser (Spectra-Physics) at 920 nm, through a 4x dry objective, 0.28 N.A. (Olympus). Scanning and image acquisition were controlled by NIS-elements software (Nikon). Brain motion artifacts interfered with imaging in awake mice but were minimized under anesthesia, enabling the stable detection of the neural activities at a single cellular resolution both in and out of the NTS area (Figure 2, Figure S3 and Supplementary Movie 1 and 2). Time-lapse imaging at various depths was performed by adjusting the imaging focal plane using a piezo to move up and down in depth (Figure 2A and Figure S3). Volumetric live imaging (2 mm x 2 mm horizontally, or smaller for zooming, 200–480 μm dorsoventrally, at around 0.4–1.0 Hz temporal resolution) of neural activities in the wide brainstem area (including the NTS) of the head-fixed mouse was performed through the implanted double prism. Importantly, this wide field of view and piezo-based volumetric imaging allowed us to detect the GCaMP fluorescence and neural activities not only from inside but also from outside of the NTS areas and confirm the specificity of the stimulus-evoked responses in the NTS, as explained below.

### Vagus nerve stimulation during NTS imaging

For the NTS imaging experiments with vagus nerve stimulation (VNS), a bipolar platinum hook electrode, connected to a programmable stimulator (SEN-8203, NIHON KODEN, Tokyo, Japan) and isolator (SS-202J, NIHON KODEN), was wrapped around the cervical vagal trunk, and was fixed and insulated from surrounding tissue using the quick-setting silicone adhesive Kwik-Sil (World Precision Instruments, FL, USA). After separating the vagus nerve from the carotid artery, a bipolar platinum hook electrode was wrapped around the left vagus nerve and fixed in place without contacting other tissues using the insulating silicone adhesive, Kwik-Sil^38,39^. The electrode was positioned to keep the positive and negative poles apart. This setup allows for stable activation of the vagus nerve while preventing irritation to the surrounding tissue. To create an anodal block for the efferent direction, the cathode and anode were located on the rostral and caudal sides, respectively, of the vagus nerve. The mechanism of the anodal block is proposed to selectively bias activation toward vagal afferent fibers^40–44^. We selectively targeted the left vagus nerve, which is usually chosen for safety reasons, as stimulation of the right vagus nerve has a greater impact on heart rate^45^. We conducted *in vivo* NTS imaging experiments after closing the surgical wound.

The mouse’s head was fixed under the objective lens by using the head plate tightly attached to the skull under anesthesia (a ketamine [70 mg/kg] and xylazine [10.5 mg/kg] mixture). VNS was triggered using custom Labview software. To determine optimal stimulation parameters VNS was delivered in blocked trials for 10 s at the frequencies of 1.0, 2.5, 5.0, 10, or 15 Hz, at the intensities of 0.1, 0.25, 0.5, or 1.0 mA, with a pulse width of 100 μsec. The maximum current intensity and frequency of stimulation were set at 1.0 mA and 10 Hz according to the previous study^37,47^. The first VNS train was delivered at least 10 s after the start of imaging, and VNS was repeated three times in total (Figure 3 and Figures S4, S5), or more (Figure 2), with a 50-sec interval. Neither facial nor muscle twitching (an indication of off-target stimulation of musculature surrounding the vagus nerve) was observed with any parameters used in any of the mice included in the present study. VNS and NTS imaging were performed with the mouse head fixed under the microscope objective lens using the head plate. The timings of the imaging and VNS were synchronously recorded with image acquisition (GCaMP imaging in NTS) using NI software (Labview; National Instruments) and NI-DAQ (National Instruments).

### Intravenous administration of CCK during NTS imaging

To examine the neural activity change in response to more physiological stimulation than VNS, we employed cholecystokinin octapeptide (CCK); it has been demonstrated that intravenous administration of CCK activates vagal afferents and increases firing rate of hepatic branch of the vagus nerve^53^ (Cox and Randich, 1997). On the day of the experiment, mouse’s head was fixed under the objective lens by using the head plate tightly attached to the skull under anesthesia (1% isoflurane). During the Ca^2+^ imaging, CCK solution (2.0 μg/kg) or saline was mixed with sulforhodamine 101 (SR101; 0.1 mM) and intravenously administered through the tail vein. 60 sec after the start of the imaging, saline with SR101 was intravenously administered and performed for 10 minutes. Using the same animal, 60 sec after the beginning of the imaging, CCK with SR101 was intravenously administered and performed for 15 minutes. Intravenous administration and NTS imaging were performed with the mouse head fixed under the microscope objective lens using the head plate.

### Imaging data analyses and statistics

The activity of NTS neurons was recorded by imaging fluorescence changes with a multiphoton microscope (Nikon A1R MP, Nikon). Time-lapse imaging at various depths was performed by adjusting the imaging focal plane using a piezo to move up and down in depth (Figure 2A and Figure S3). The volumetric live imaging (at around 0.4–1.0 Hz temporal resolution) of NTS neural activities was performed through the implanted double prism. We recorded a total of 10–16 focal planes (z-planes) at each time point for each mouse. In an example case (Figure 2D), we recorded a total of 16 z-planes at each time point. We stacked every 4 consecutive plane from top to bottom to create an average image for each time point. This stacking was done with a two-plane overlap; for instance, the first stack included the 1st to 4th z-planes, the second stack included the 3rd to 6th z-planes, and subsequent stacks followed this pattern. As a result, we obtained 7 stacked planes in total in this experiment, labeled as Zp1 through Zp7, respectively. This procedure is advantageous to enhance the signal to noise ratio in each movie, which is critical for further automatic detection of the active neurons^48^. After the automatic detection of the active neurons (details for this detection method are described below), we checked the existence of the overlapped detection of the regions of interest (ROIs, i.e., active neurons) between adjacent Zps next each other, and removed such overlaps by the manual inspection.

Importantly, before stacking 4 consecutive z-planes at each time point as explained above, they were processed to correct for brain motion artifacts using the enhanced correlation coefficient image alignment algorithm^62^. For this z-axis correction, first (most dorsal surface frame) was used as a reference for correcting the second z-plane. Then, an average image of these two aligned z-planes was further used for the correction of the third z-plane. Eventually an average image of these three aligned z-planes was used for the motion correction of the fourth z-plane. Through this procedure, four z-planes were motion corrected and used to accurately obtain a stacked (averaged) image for each Zp at each time point. Next, the time-series of stacked GCaMP images at each Zp was further processed to correct for brain motion artifacts using the enhanced correlation coefficient image alignment algorithm^62^. Temporal dynamics of neural activity were clearly observed at multiple depths after this motion correction procedure (Supplementary Movies 1 and 2).

These motion-corrected movies were used for automatic ROI detection. The ROIs for the detection of neural activity were automatically selected using a constrained nonnegative matrix factorization algorithm in MATLAB R2014a/R2019b as described previously^48,49^, with some manual adjustments as described above. This detection method is advantageous in isolating the activity of the individual neuron, even when other active neurons are nearby or overlapped in the field of view^48^. The same constrained nonnegative matrix factorization package for ROI detection also provides an option for signal processing and measurement of the signal change (ΔF/F) of each neuron. We used this method to identify neurons and calculate ΔF/F for them. While neurons for the analyses were initially automatically detected, neurons responding to noisy signals with no apparent calcium transient at any time during the experimental days were identified by visual inspection and excluded from further analysis. To apply the same regions of interest (ROIs) for analyzing images obtained across multiple conditions (i.e., various VNS conditions or saline vs CCK experiments), the movies from the same mouse and the same field of view were precisely aligned with each other using the same enhanced correlation coefficient algorithm as above, and these multiple movies of the same field of view were tentatively concatenated for the ΔF/F measurement. The obtained ΔF/F data were eventually separated to evaluate the neural responses under each condition.

For the statistical analysis, we used MATLAB R2022a/R2023b. The Wilcoxon signed-rank test for a paired comparison and the Kolmogorov-Smirnov test for a two-group comparison with pooled data (from all neurons) were used to determine statistical significance (*p* < 0.05) in Figure 4. Two-tailed tests were selected for all statistical analyses. Graphs were produced by MATLAB R2022a/R2023b.

As for the imaging data analysis, one concern was that the off kinetics of the GcaMP6f signal might not be sufficiently fast to consider the responses to 3 trails of VNS as independent data, and thus statistical evaluation based on these responses might not be reliable to evaluate the neural responses. As shown in Fig. 3G, however, neurons showed a reduced response to repeated stimulation, indicating that calcium signaling was not enhanced by delayed off-kinetics.

After the CCK administration, we observed prolonged neural activation, which was detected as continuous calcium transients (Figure 4 and Figure S6). To evaluate the event frequency of the transients, we identified the timing of the calcium transients by analyzing the first derivative of ΔF/F (Figure S6). The first derivatives of each neuron, from the entire imaging session (including both the saline and CCK experiments), were thresholded at 0.5 times the standard deviation to detect active time points. The onset of each event was then used to determine the timing of each transient. The intervals between calcium transient events were then calculated and used to compare the event frequencies between the saline and CCK experiments (Figure 4F).

### Drugs

CCK was purchased from PEPTIDE INSTITUTE, INC. and was dissolved in saline (2.0 μg/kg). In addition, sulforhodamine (SR101), a red fluorescent dye, is diluted to 0.1 mM in saline or CCK solution to confirm the successful intravenous administration and the injection timing during NTS imaging.

## Supplementary Materials

Supplementary Materials (Figures. S1 to S6, and caption for Supplementary Movies S1 to S5) are explained in a separate document.

