## Supplementary Information for "Low-invasive, wide-field, and cellular resolution two-photon imaging of neural population activity in brainstem and nucleus tractus solitarii"

**This PDF file includes:**

Figures. S1 to S6

Captions for Movies S1 and S5

**Other Supplementary Materials for this manuscript include the following:**

Movies S1 and S5

#### Figure. S1

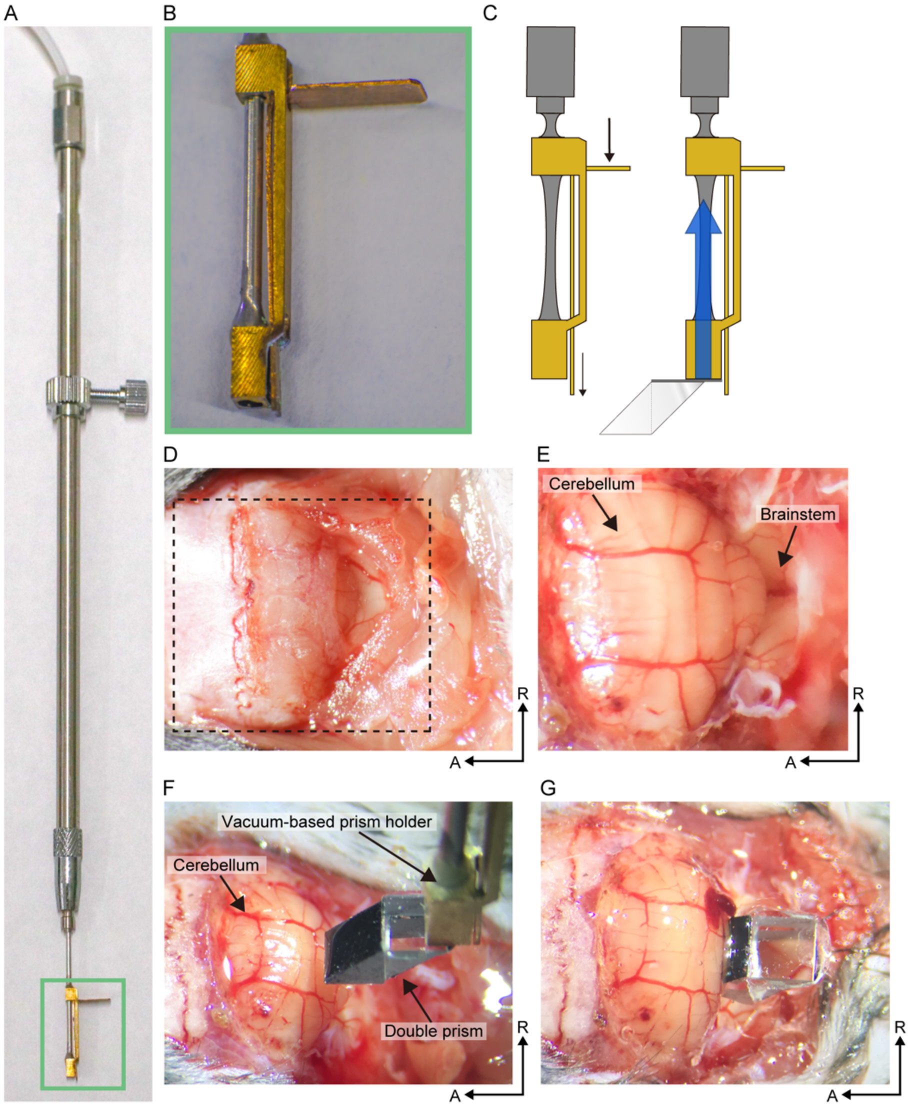


**Supplementary Figure 1.** A vacuum-based prism holder and implantation of the double prism

(A) Image of the vacuum-type prism holder, which vacuums the double prism while implanting it between the cerebellum and brainstem.

(B) Magnified view of the vacuum-type prism holder (indicated by the green line in panel A).

(C) Schematic diagram showing how to move the holder's hook (left) and how to vacuum the double prism assembly (right). The thick black arrow shows the direction in which the hook is pushed. The thin black arrow indicates the hook's movement when pushed. The blue arrow indicates the direction of vacuuming.

(D) Image before removing the skull and dura over the cerebellum. The dotted line indicates the area where the skull will be removed (as shown in D).

(E) Image after removing the skull and the dura mater covering the cerebellum and brainstem.

(F) Image of the vacuum-type prism holder attaching the prism before implanting it between the cerebellum and brainstem.

(G) Image after inserting the double prism between the cerebellum and brainstem. UV craft resin was used to fix the double prism and coat the surface of the targeted nerve structures. After the UV resin had cured, the holder was carefully released for the further head plate attachment (Figure 1F).

A, anterior; R, right.

### Figure. S2

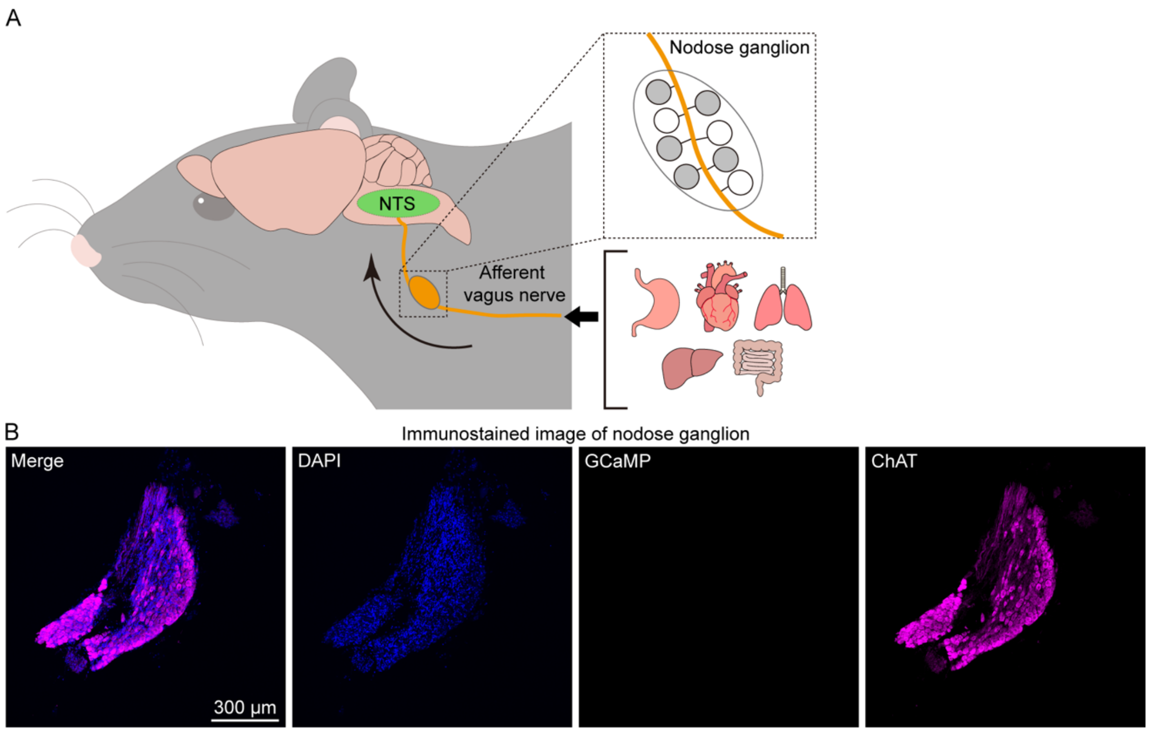


**Supplementary Figure 2.** No expression of GCaMP in nodose ganglion neurons

(A) Schematic illustration of afferent vagal projections from visceral organs to the NTS.

(B) Immunohistochemical staining confirmed no expression of GCaMP (green) in the nodose ganglia, confirming that the afferent vagal fibers terminating at the NTS area did not express GCaMP. DAPI (blue) and ChAT (magenta).

#### Figure. S3

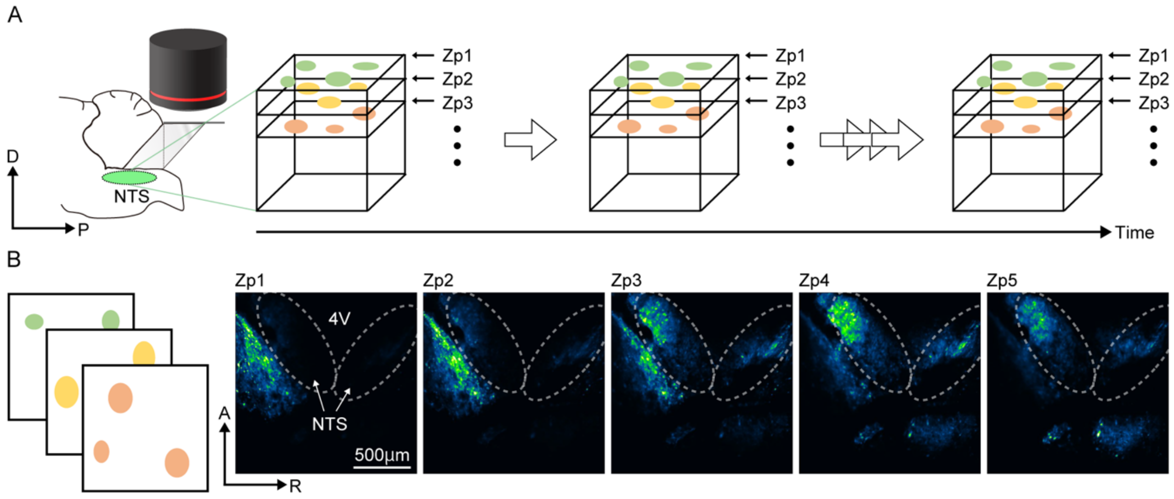


**Supplementary Figure 3.** Volumetric time-lapse imaging of brainstem neural activity

(A) Two-photon imaging was performed through the objective lens (black) and the implanted double prism in the head-fixed mouse under anesthesia. Time-lapse imaging at various depths was performed by adjusting the imaging focal plane using a piezo to move up and down in depth. Colored circles illustrate examples of detected regions of interest (ROIs) at Zp1, Zp2, and Zp3.

(B) Examples of *in vivo* volumetric live imaging (at most 2 mm x 2 mm horizontally, at around 0.5–1.0 Hz temporal resolution) of neural activities in the wide brainstem area (including the NTS). The results from the same mouse are also shown in Figures 1G, 2D and Supplementary Movie 1. Using the wide field of view and horizontal coordinates, the position of the fourth ventricle serves as a key reference for locating the NTS. In this example case, a total of 16 z-planes at each time point were measured and stacked every 4 consecutive planes from top to bottom to create average images (termed Zp, z-stacked horizontal plane) for each time point. This stacking was done with a two-plane overlap; for instance, the first stack included the 1st to 4th z-planes, the second stack included the 3rd to 6th z-planes, and subsequent stacks followed this pattern. As a result, we obtained 7 stacked planes in total in this experiment, labeled as Zp1 through Zp7 respectively. Here, Zp1 to Zp5 are shown (Zp1, 2 and 4 are the same data as Figure 2D). In this panel, median fluorescence over the entire imaging period for each Zp is presented in green-fire-blue to show the baseline fluorescence level. See Methods for details.

D, dorsal; P, posterior; A, anterior; R, right.

### Figure. S4

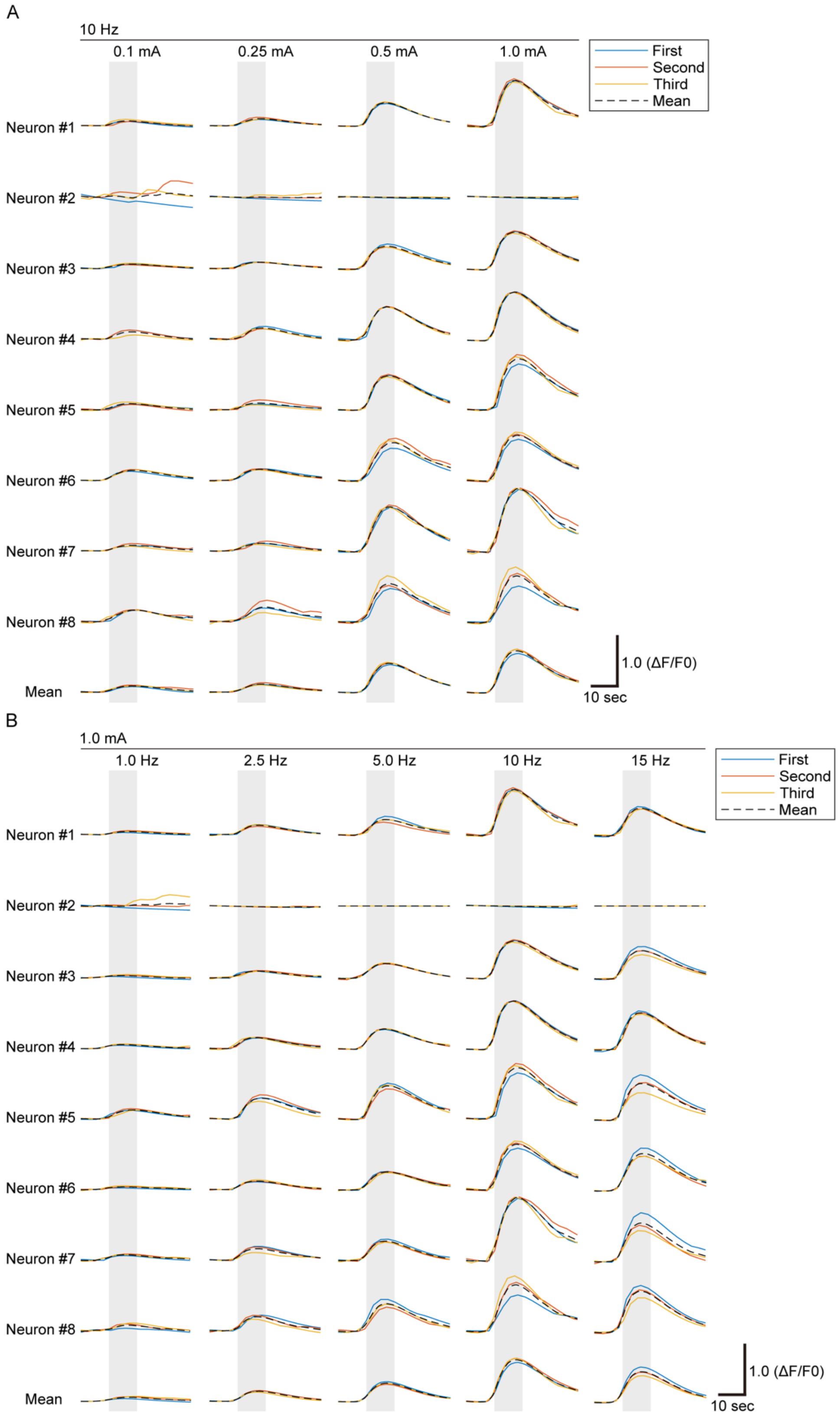
A
10 Hz
0.1 mA
0.25 mA
0.5 mA
1.0 mA
First
Second
Third
Mean
Neuron #1
Neuron #2
Neuron #3
Neuron #4
Neuron #5
Neuron #6
Neuron #7
Neuron #8
Mean
1.0 (ΔF/F0)
10 sec
B
1.0 mA
1.0 Hz
2.5 Hz
5.0 Hz
10 Hz
15 Hz
First
Second
Third
Mean
Neuron #1
Neuron #2
Neuron #3
Neuron #4
Neuron #5
Neuron #6
Neuron #7
Neuron #8
Mean
1.0 (ΔF/F0)
10 sec

**Supplementary Figure 4.** Raw neural activity data of recorded NTS neurons in response to various VNS conditions

(A) Change in GCaMP signal (ΔF/F0) of each neuron in the frequency-fixed (10 Hz) condition with various intensity of 0.1, 0.25, 0.5, and 1.0 mA. The 10-sec VNS, indicated by gray shaded area, was delivered three times at 50-sec interval (same as used in Figure 3). Colored lines indicate the neural responses to the first, second, and third VNS. Dashed line indicates mean of the responses to three VNSs. These data are summarized in Figure 3.

(B) Change in GCaMP signal (ΔF/F0) of each neuron in the intensity-fixed (1.0 mA) condition with various frequencies of 1.0, 2.5, 5.0, 10, and 15 Hz.

### Figure. S5

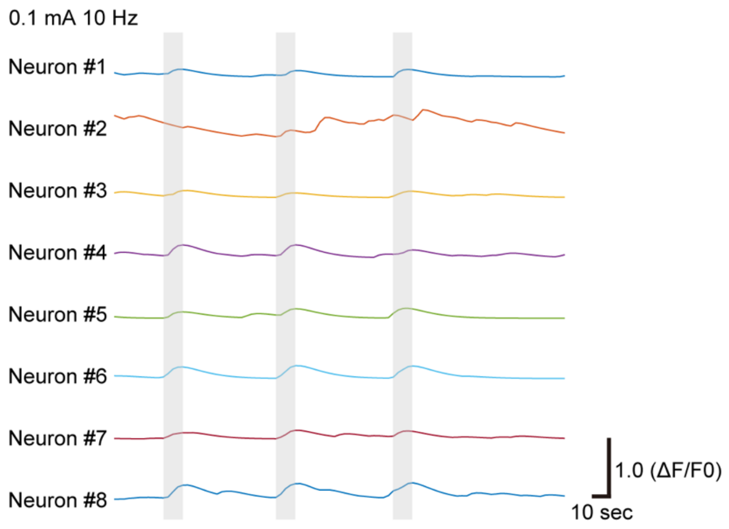


**Supplementary Figure 5.** Neural responses of NTS neurons induced by minimal intensity VNS

Changes in GCaMP signals (ΔF/F0) of NTS neurons were detected (in neurons #1, #3–#8) following all three VNS at the minimal intensity of 0.1 mA and frequency of 10 Hz. On the other hand, one neuron (neuron #2) located outside of the NTS showed only spontaneous activity change and no clear response to the VNS.

### Figure. S6

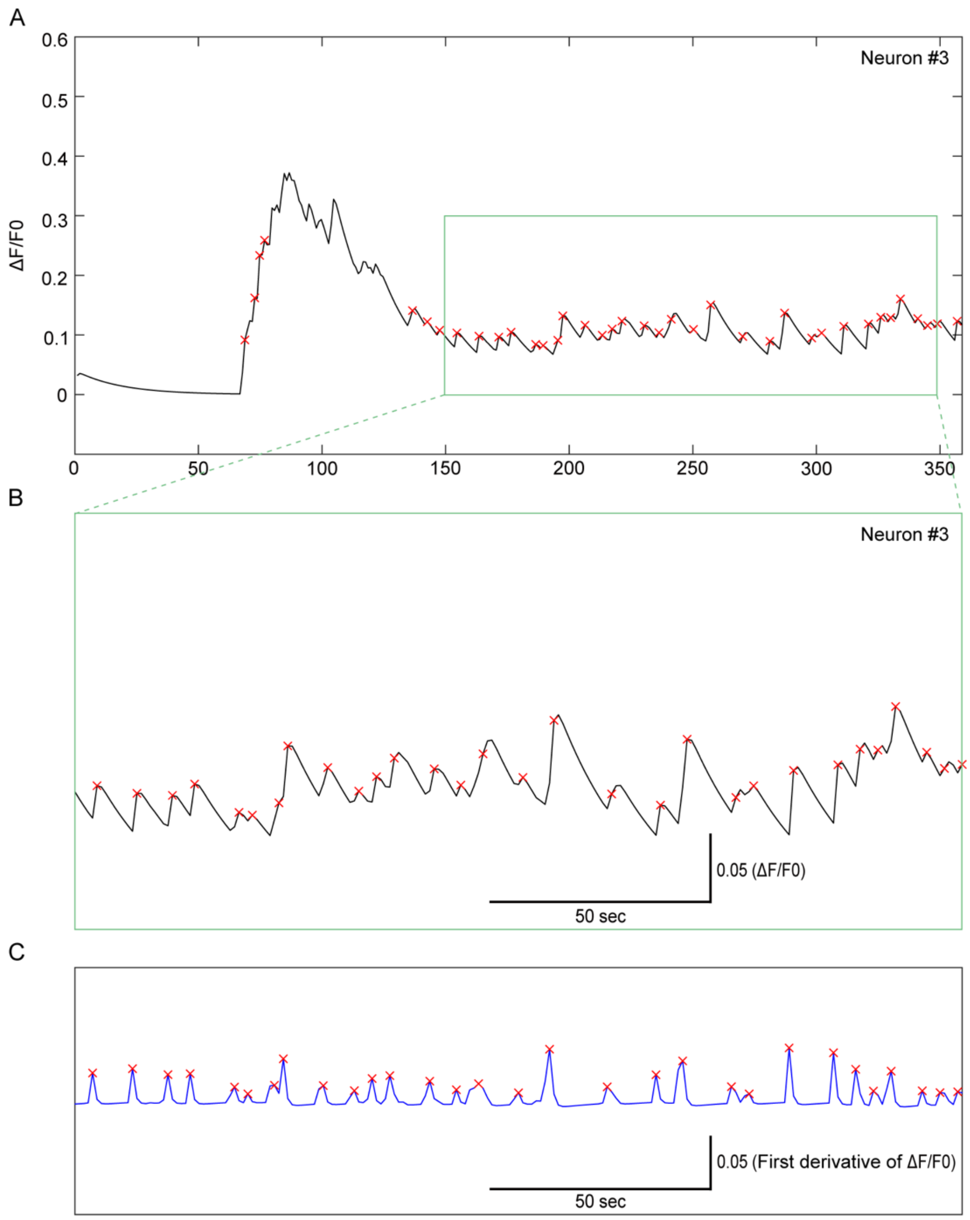


**Supplementary Figure 6.** Detection of prolonged neural activation in NTS neurons after CCK administration

(A) Changes in GCaMP signals (ΔF/F0) after CCK administration. A large signal immediately after CCK administration was followed by smaller, continuous calcium transients, indicating prolonged activation of neurons in the NTS. (B) Magnified view of a section from (A). (C) First derivative of ΔF/F0. To identify the timing of the calcium transients, the first derivative of ΔF/F0 was analyzed. The detected transients are marked with red crosses.

#### Supplementary Movies

**Supplementary Movie 1.** An example of neural activities in the brainstem around the NTS and their responses to VNS, detected by changes in GCaMP signals. AAVdj/Syn.GCaMP7f was used for this experiment. The results from the same mouse are also shown in Figure 2D and S3. Time-lapse imaging at various depths was performed by adjusting the imaging focal plane using a piezo. Volumetric live imaging (at approximately 0.4–1.0 Hz temporal resolution) of NTS neural activities was performed through the implanted double prism. This movie shows images from Zp1 to Zp5. Image at each Zp was processed with a median filter of radius 1 pixel applied along the time axis. These five movies were horizontally combined. The images were color-coded using the green-fire-blue lookup table in ImageJ according to the fluorescence level of the GCaMP signal. During the VNS, the character "VNS" is displayed in the top left corner of the movie.

**Supplementary Movie 2.** High-magnification imaging of NTS neural activities in response to VNS. AAV1/Syn.GCaMP6f was used for this experiment. The results from the same mouse are also shown in Figure 2E. This movie shows images from Zp1 and Zp5. Image at each Zp was processed with a median filter of radius 1 pixel applied along the time axis. These movies were horizontally combined. During the VNS, the character "VNS" is displayed in the top left corner of the movie.

**Supplementary Movie 3.** Visualization of the timing of intravenous administration of saline during *in vivo* NTS imaging. Time 0 sec indicates the timing of the intravenous administration of a saline solution mixed with the fluorescent dye SR101 (magenta) via the tail vein. By monitoring the SR101 fluorescence, we confirmed the successful intravenous administration. Neural activities of the same experiment (saline administration) is also shown in Supplementary Movie 4.

**Supplementary Movie 4.** Almost no neural responses to saline intravenous administration monitored by changes in GCaMP fluorescence in the brainstem around the NTS area. AAVdj/Syn.GCaMP7f was used for this experiment. The results from the same mouse are also shown in Figure 4. Time 0 sec shown in the top right corner of the movie indicates the timing of saline administration. The baseline-subtracted signal is shown in green over a background of the baseline fluorescence image in gray.

**Supplementary Movie 5.** Neural activities in the brainstem around the NTS area in response to intravenous administration of CCK detected by changes in GCaMP fluorescence. AAVdj/Syn.GCaMP7f was used for this experiment. The results from the same mouse with saline are shown in Supplementary Movie 4. Time 0 sec shown in the top right corner of the movie indicates the timing of CCK administration. The baseline-subtracted signal is shown in green over a background of the baseline image in gray.
